# Genomic Evidence of Multidrug-Resistant *Salmonella* in Wild Waterbirds from High-Andean Lakes of Ecuador

**DOI:** 10.64898/2026.04.09.717384

**Authors:** Nathaly Reyes, Christian Vinueza-Burgos, José Medina-Santana, María L. Ishida, Brian D. Sauders, Deysi Anchatuña, Nivia Luzuriaga-Neira

## Abstract

*Salmonella* spp. represents a leading cause of foodborne disease globally. Wild aquatic birds inhabiting ecosystems impacted by human activities may serve as reservoirs and dispersers of *Salmonella* and antimicrobial resistance genes (ARGs), posing significant public health risks. This study evaluated the prevalence, serovars, resistance genes, and genomic relationships of *Salmonella* in fecal samples from wild aquatic birds across three high-Andean lakes in Ecuador. Of 134 samples collected from 10 species, five (3.73%) tested positive, all from Yahuarcocha Lake, isolated from *Fulica ardesiaca* and *Phalacrocorax brasilianus*. Two serovars were identified: *Salmonella* Infantis (ST32, n=4) and *Salmonella* Newport (ST45, n=1). Three *S.* Infantis isolates exhibited multidrug resistance (MDR), mediated by a pESI-like plasmid carrying resistance genes against beta-lactams, aminoglycosides, tetracyclines, sulfonamides, trimethoprim, fosfomycin, and chloramphenicol. SNP-based phylogenetic analysis revealed low genetic divergence (≤10 SNPs) between wildlife and poultry-associated isolates, indicating a shared transmission network. These findings support a likely spillover from poultry production systems into wild bird populations, and highlight the role of wild aquatic birds as ecological sentinels and potential disseminators of MDR *Salmonella* across interconnected human, animal, and environmental systems. These results underscore the need to incorporate human, animal, and environmental health factors within a One Health framework.

## Introduction

*Salmonella*, a genus of Gram-negative bacteria, represents a major public health concern worldwide due to its role as one of the leading causes of foodborne diseases [1,2]. In the United States alone, foodborne salmonellosis results in economic losses of approximately 3 billion U.S.D. per year and is associated with about 420 deaths each year [3].

The ecological dynamics of *Salmonella* extend beyond agricultural environments and include diverse natural ecosystems inhabited by wildlife. Wild birds feeding in anthropogenically impacted habitats, including lakes, rivers, and coastal environments, may become exposed to environmental sources of this pathogen [4]. Human activities, including urbanization, agriculture, livestock production, and industrial development, create ecological interfaces that facilitate interactions among wildlife, domestic animals, and humans, potentially increasing opportunities for pathogen transmission [5].

Although food-producing animals represent a major source of Salmonella contamination, free-living and migratory birds have also been recognized as reservoirs of diverse *Salmonella* serovars [6–9]. Waterbirds, seabirds, and shorebirds inhabit a wide range of ecosystems, including freshwater lakes, rivers, wetlands, and coastal environments. Species such as the Neotropical Cormorant (*Phalacrocorax brasilianus*), Double-crested Cormorant (*Phalacrocorax auritus*), and Dominican Gull (*Larus dominicanus*) have been documented as reservoirs for different serotypes of *Salmonella* [6,10]. Infections have also been documented in shorebirds, passerines, and other aquatic bird species [11–13].

The interaction between Salmonella and wild birds is influenced by multiple ecological factors, including habitat characteristics, migratory behavior, interspecific interactions, and anthropogenic environmental pressures [14,15]. Wild birds may acquire *Salmonella* through contaminated water sources, fecal–oral transmission, or contact with infected wildlife or domestic animals [16–18]. Although infections in wild birds are frequently asymptomatic, clinical disease may occur, particularly in juvenile individuals (The Center for Food Security and Public Healt 2006; Uribe and Suárez 2006).

Wild birds also serve as valuable ecological indicators of environmental health despite their ability to respond rapidly to environmental change. Aquatic birds are particularly sensitive to fluctuations in food availability, habitat quality, and environmental contamination. Consequently, they are frequently used as bio-indicators of ecosystem disturbance and environmental pollution [21]. Previous studies have shown associations between habitat disturbance and increased prevalence of infectious diseases such as avian influenza, vector-borne diseases, and bacteria carrying antimicrobial resistance genes [22,23].

Because many bird species migrate long distances, they may contribute to the geographic dissemination of antimicrobial-resistant bacteria. Antimicrobial resistance genes can persist in the environment and spread within microbial communities through horizontal gene transfer, thereby facilitating their propagation and increasing the risk of environmental dispersion [24–26].

Therefore, this study aimed to evaluate the occurrence of *Salmonella* in wild aquatic birds inhabiting three high-Andean lakes in Ecuador and to characterize the detected strains through phenotypic and genomic analyses. Specifically, we assessed the presence, serotypes, antimicrobial resistance profiles, resistance genes, and clonal relationships of *Salmonella* isolates obtained from fecal samples collected from wild birds in the Andean lakes Yahuarcocha, Yambo, and Colta.

## Material and Methods

### Study sites

The study was conducted in three high-Andean lakes of Ecuador: Yahuarcocha (00°22’N and 78°06’W), Yambo (1°6’0’S and 78°34’60’W), and Colta (1°43’60’S and 78°45’0”). These lakes were selected because they represent important aquatic ecosystems in the northern and central Andes, and support diverse avian communities that include both endemic and seasonally migratory species with distinct ecological behaviors and feeding strategies [27]. The lakes are distributed along an altitudinal gradient from approximately 2,200 to 3,300 m *a.s.l.*, with Yahuarcocha situated at 2,210 m, Yambo at 2,560 m, and Colta at 3,310 m. Despite this gradient, the sites share broadly comparable ecological conditions characterized by cool Andean climates with mean annual temperatures from ∼8-18 °C and relative humidity typically between 60% and 80% [28]. These wetlands support aquatic vegetation and invertebrate communities that provide feeding resources for waterbirds.

The three lakes exhibit varying degrees of eutrophication, driven by nutrient inputs from surrounding agricultural and human activities. However, this condition is particularly pronounced in Yahuarcocha Lake, where urban development, tourism, and agricultural land use contribute to increased nutrient loads and changes in water quality within the ecosystem [29]. Field sampling was conducted under the legal permit MAE-CGZ1-DPAI-2017-1524-O issued by the Ecuadorian environmental authority.

### Field Sampling

Sampling was conducted during two periods corresponding to seasonal bird migration, in October 2017 and March 2018. Birds were captured at night from a motorboat using spotlight distraction techniques. Captured individuals were immediately placed in ornithological bags and transported to a field laboratory near each study site, where they underwent species identification, physical examination, and individual coding [28]. Fecal samples were collected from birds while temporarily held in the bags. All birds were released at their capture site within approximately 10 min.

A total of 134 fecal samples, each weighing at least 3 g, were collected from 10 distinct species of resident and migratory waterbirds (Table 1). Samples were coded, stored, and transported to the laboratory under refrigerated conditions (4°C) for subsequent analysis. All procedures were performed by trained personnel following standard animal handling guidelines to minimize stress [30]. No birds were harmed or injured during capture, sampling, or release.

**Table 1.** Prevalence of *Salmonella* spp. in fecal samples from wild aquatic birds across three high-Andean lakes of Ecuador (October 2017 – March 2018). Andean lakes of Ecuador.

|  |  | Andean lakes of Ecuador |  |  |  |
| --- | --- | --- | --- | --- | --- |
|  |  | Yahuarcocha | Yambo | Colta | Total % |
|  |  | % (n/N) | % (n/N) | % (n/N) | (n/N) |
| Bird species |  |  |  |  |  |
| Scientific name | Common name |  |  |  |  |
| *Fulica ardesiaca* | Andean coot | 21.0 (4/19)<sup>a,b</sup> | 0 (0/23) | 0 (0/6) | 8.3 (4/48) |
| *Phalacrocorax brasilianus* | Neotropical Cormorant | 5.9 (1/17)<sup>a</sup> | - | 0 (0/1) | 5.5 (1/18) |
| *Ardea alba* | Snowy egret | 0 (0/4) | - | - | - |
| *Gallinula chloropus* | Swamp chicken | 0 (0/6) | - | - | - |
| *Podiceps occipitalis* | Silvery Grebe | 0 (0/6) | - | 0(0/10) | - |
| *Oxyura ferruginea* | Andean duck | 0 (0/5) | 0 (0/5) | 0(0/16) | - |
| *Anas georgica* | Yellow-billed Pintail | 0 (0/1) | 0 (0/10) | - | - |
| *Bubulcus ibis* | Cattle egret | 0 (0/1) | - | - | - |
| *Anas andium* | Andean teal | - | 0 (0/3) | - | - |
| *Anas discors* | The blue-winged teal | - | 0 (0/1) | - | - |
| **Total** |  | **8.4 (5/59)** | **0 (0/42)** | **0 (0/33)** | **3.7 (5/134)** |

**Notes:** Positive samples : a: *S. Infantis*, b: *S. Newport*

**Table 2.**
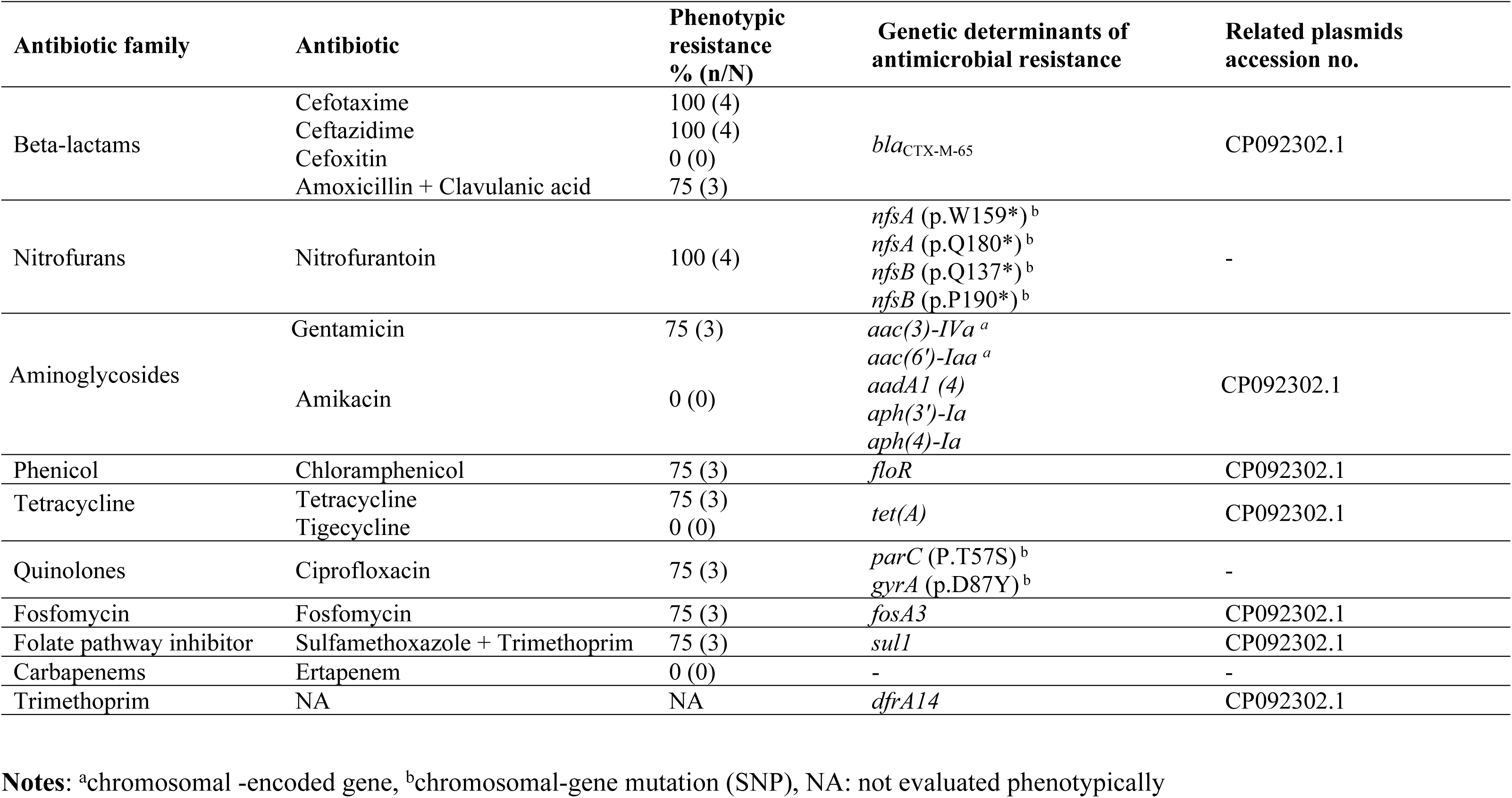
Phenotypic antimicrobial resistance and associated genetic determinants identified in Salmonella Infantis isolates from wild aquatic birds at Yahuarcocha Lake, Ecuador.

### Ethics statement

Wild birds were captured and handled for fecal sample collection under permits MAE-CGZ1-DPAI-2017-1524-O (local permit) and MAE-DNB-CM-2019-0119 (national permit) issued by the Ecuadorian environmental authority (Ministerio del Ambiente del Ecuador). At the time of sampling, no institutional animal ethics committee was available; therefore, authorization was granted by the national environmental authority under the above-mentioned permits.

### Microbiological analysis, DNA extraction, and sequencing

*Salmonella* isolation and identification were performed following the ISO 6579-1:2017 standard protocol. Antimicrobial susceptibility testing of isolates was conducted using the Kirby-Bauer disk diffusion method, according to the Clinical and Laboratory Standards Institute guideline [31]. For phenotypic resistance determination, bacterial suspensions adjusted to a 0.5 McFarland standard were inoculated into Mueller-Hinton agar. After incubation at 37 °C for 18–24 h, inhibition zone diameters were measured and interpreted according to CLSI criteria; intermediate susceptibility values were considered resistant. The antibiotics tested included Cefotaxime (CTX, 30 µg), Ceftazidime (CAZ, 30 µg), Cefoxitin (FOX, 30 µg), Amoxicillin + Clavulanic acid (AMC, 30 µg), Nitrofurantoin (F, 300 µg), Gentamicin (CN, 10 µg), Amikacin (AK, 30 µg), Chloramphenicol (C, 30 µg), Fosfomycin (FF, 200 µg), Sulfamethoxazole + Trimethoprim (SXT, 25 µg) and Ertapenem (ETP, 10 µg), and Ciprofloxacin (CIP, 5 µg).

DNA extraction was performed using the Wizard SV Genomic DNA Purification System kit from Promega (Wisconsin, United States), following the manufacturer’s instructions. DNA quantity and quality were assessed using the Quantum Fluor System and Nanodrop. Samples with a minimum concentration of 10 ng/μL and an OD 260/280 ratio between 1.75 and 2.05 were used for subsequent sequencing applications.

### Bio-informatic analysis

The genomic analysis followed the methodology previously described by Medina-Santana *et al*. (2022). Briefly, raw sequencing reads were processed through EnteroBase for quality control, trimming, and assembly, as described by Achtman *et al.* [33] and Alikhan *et al*. [34]. EnteroBase was utilized for serotype identification, sequence type (ST) identification via multi-locus sequence typing (MLST), core genome MLST (cgMLST) analysis, and SNP-based phylogenetic tree construction. The assemblies were integrated into the downstream analytical pipeline.

To assess the population structure and genetic relatedness among isolates from different sources, Principal Components Analysis (PCA) and graphical visualizations were performed in R [35] using the *adegenet* package [36,37]. Phylogenetic analyses and tree visualization were conducted using the packages: *ape*, *phangorn*, and *ggtree*, with additional graphical customization and data manipulation performed using: *ggplot2*, *ggnewscale*, and *dplyr*. Metadata and the complete list of genomes included in the comparative analysis are provided in Supplementary Information (Table S1). All scripts and associated data are available at the project repository:https://github.com/Nivia-L/Salmonella_wild_birds.git.

SNP tree analysis was visualized and annotated using iTOL v. 6.0, following the method described by Letunic and Bork (2021). Genomic comparisons were performed among isolates from this study and others sourced from various locations within Ecuador, all data accessible through the Enterobase database [38]. Genes and single-nucleotide polymorphisms (SNPs) associated with antimicrobial and disinfectant resistance were identified using the AMRfinder plus and ResFinder databases, as detailed by Zankari *et al*. [39] and Feldgarden *et al*.[40]. Virulence genes were identified from the Virulence Factor Database (VFDB) using a mass screening of contigs in ABRicate software following the methodology outlined by Chen *et al*. [41], and Seemann [42].

Functional nitrofurantoin resistance-associated mutations were identified following Medina-Santana *et al*. (2022). All genomes were aligned to the wild-type of nfsA and nfsB genes, which encode oxygen-insensitive nitroreductase enzymes in *S.* enterica (accession number NC_003197), using Geneious Prime version 2021.0.3. Subsequently, sequences harboring mutations were subjected to in silico translation and aligned with the wild-type protein sequences to visualize nonsense mutations.

Plasmid identification was performed using Platon v.1.6 [43], which predicts plasmid contigs from assembled sequences. Additionally, Geneious Prime V.2022.1.1 (https://www.geneious.com) and NCBI-BLAST [44] were employed to ascertain the identity of the plasmids. To achieve this, tentative plasmids were mapped across all genomes using the map-to-reference tool within Geneious Prime software.

## Results

*Salmonella* was detected in five of the 134 samples (3.73%; 95% CI: 0.5–6.9%). All positive samples were obtained from Yahuarcocha Lake (Table 1). The isolates were recovered from two avian species: the Andean Coot (*Fulica ardesiaca*), a resident species, and the Neotropical Cormorant (*Phalacrocorax brasilianus*), a migratory aquatic bird.

Among the isolates, two different serotypes were determined. Four isolates were identified as *Salmonella* Infantis, while one isolate corresponded to *Salmonella* Newport. All *S.* Infantis isolates belonged to sequence type ST32, whereas the *S*. Newport isolate was assigned to ST45. Therefore, this isolate was excluded from the dendrogram of genetic relatedness.

### Antimicrobial resistance and plasmids identification

#### Phenotypic profiles

The phenotypic profiles showed that three *S.* Infantis isolates were multidrug-resistant (MDR), characterized by resistance to three or more antibiotic classes. The predominant resistance was observed against nitrofurans (nitrofurantoin), beta-lactams (cefotaxime, ceftazidime, and clavulanic acid), aminoglycosides (gentamicin), and tetracyclines, quinolones, fosfomycin, sulfamethoxazole + trimethoprim, and chloramphenicol. In contrast MDR to carbapenems was not detected in any of the strains (Table 2). In contrast, *S.* Newport demonstrated overall susceptibility to most tested antibiotics, with resistance observed exclusively to gentamicin (an aminoglycoside) and nitrofurantoin (a nitrofuran).

#### Antimicrobial resistance genes (AMR genes)

All strains of *S.* Infantis harbored genetic determinants conferring resistance against nine antimicrobial classes (refer to Table 2), except for carbapenems. AMR gene blaCTX-M-65 was detected in all isolates, consistent with the observed phenotypic resistance to cefotaxime and ceftazidime. Genes conferring resistance to aminoglycosides included aac(3)-IVa, aac(6’)-Iaa, aadA1, aph(3’)-Ia, and aph(4)-Ia, while the floR gene was identified in isolates resistant to chloramphenicol. Tetracycline resistance was mediated by the tet(A) gene, and the fosA3 gene was associated with resistance to fosfomycin. Additionally, the sul1 gene conferred resistance to sulfonamides, and dfrA14 was linked to trimethoprim resistance. Mutations in gyrA (p.D87Y) and parC (p.T57S) were identified in quinolone-resistant strains, whereas mutations in nfsA and nfsB were related to resistance to nitrofurantoin. These resistance determinants were located on a plasmid with accession number CP092302.1, suggesting potential for horizontal gene transfer. The *S.* Newport strain harbored the chromosomal *parC* mutation (p.T57S) but did not show phenotypic resistance to the tested quinolone (ciprofloxacin). Additionally, the strain carried the previously undescribed mutations p.Q9K, p.G66A, p.T69V, and p.D103E in the *nfsB* gene.

### Plasmid Identification

The pESI-like plasmid PNUSAS047891, with GenBank accession number CP092302, was identified in all *S.* Infantis strains, and harbors several genes associated with antibiotic resistance, virulence, and resistance to disinfectants and heavy metals (Table 2, Figure 1). Since the repI1 locus identity remained undetermined, a perfect match to the plasmid Multilocus Sequence Type (pMLST) could not be assigned. However, the plasmid belongs to the IncI1 incompatibility group *S.* Newport did not harbor any known plasmid contigs.

**Figure 1.**
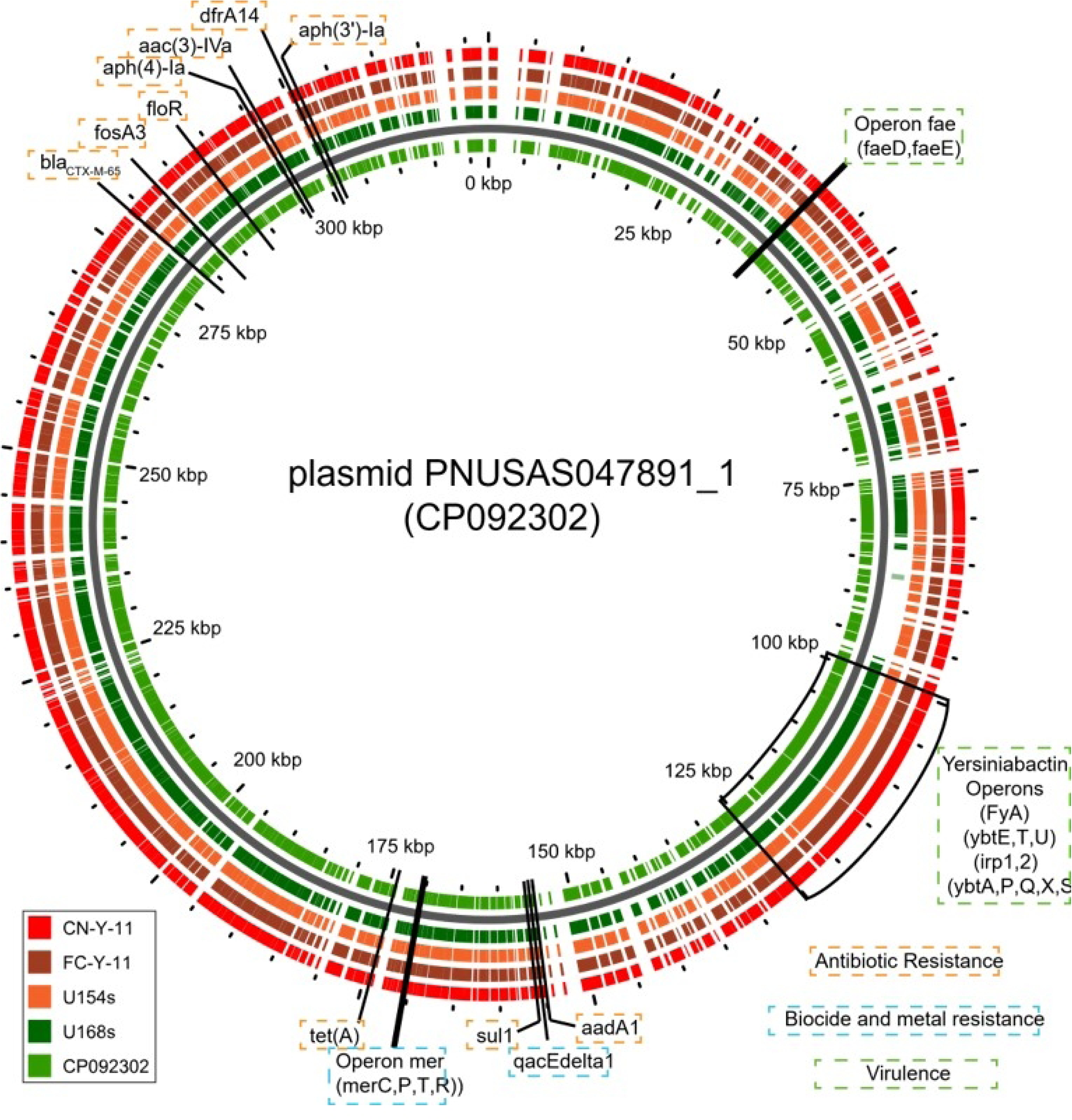
Plasmid alignment of *Salmonella* strains isolated in Yahuarcocha lake from Andean Coot (*Fulica ardesiaca*) and Neotropical Cormorant (*Phalacrocorax brasilianus*). Color-coded is related to *S.* Infantis strains belonging to avian species.

### Genotypes and Principal Components Analysis

The PCA dataset included 75 genomes obtained from EnteroBase representing multiple sources, including broilers (n = 32), layers (n = 17), pigs (n = 8), supermarket environment (n = 5), humans (n = 4), wild ducks (n = 2), and wild bird isolates generated in this study (n = 5). The distribution of isolates according to source is summarized in Supplementary Table (Table S2).

PCA based on genome-wide SNPs revealed a substantial overlap among isolates from different sources (Fig. 2). Wild bird isolates clustered within the genomic space predominantly occupied by poultry-associated strains, indicating close genetic relatedness. However, a small group of heavy-breeders formed a distinct cluster along PC1. Despite this apparent separation, statistical analysis showed no significant association between PC1 scores and isolate source (*ANOVA, F = 0.165, p = 0.919*), suggesting limited genomic differentiation among sources.

**Figure 2.**
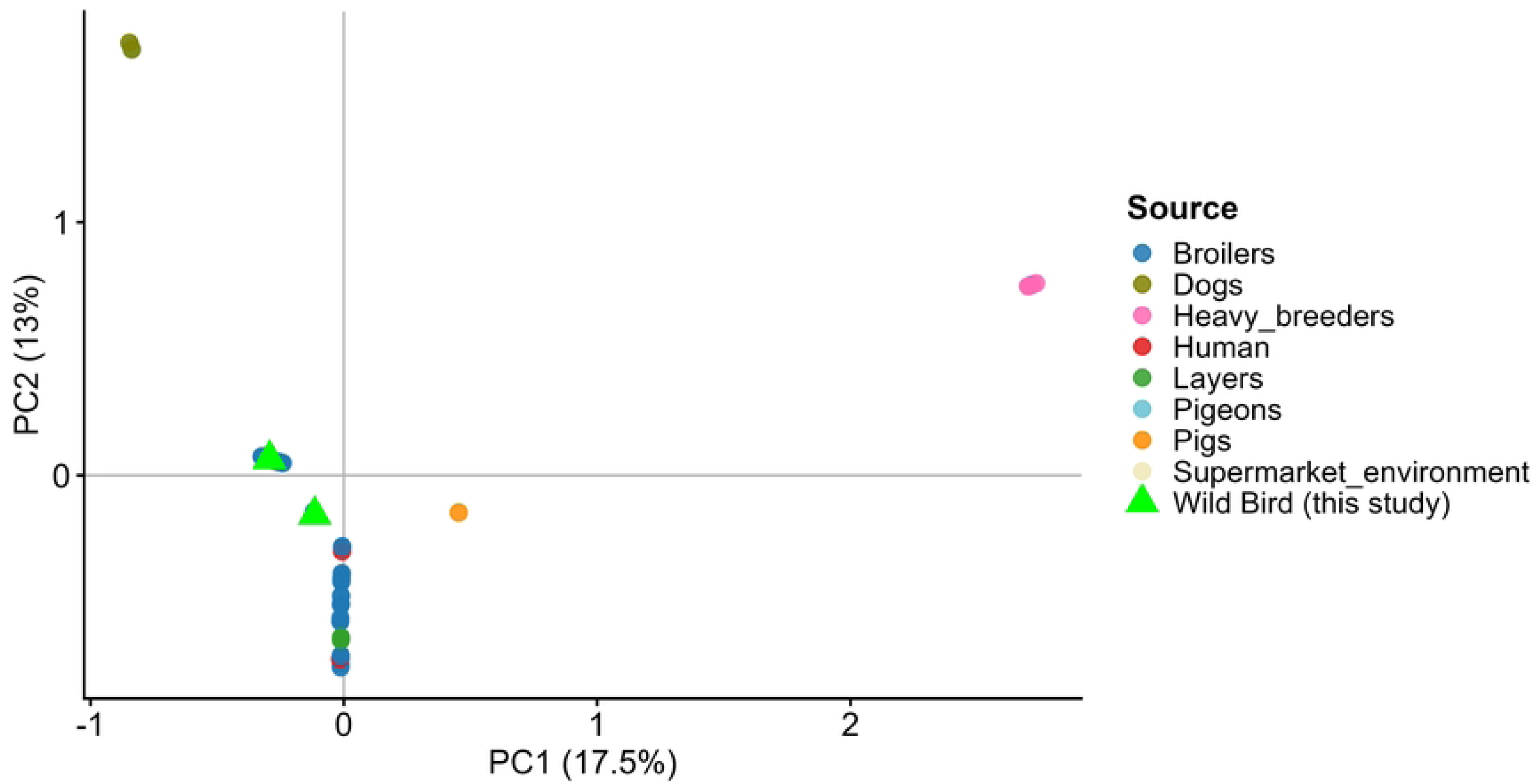
Principal Component Analysis (PCA) of genome-wide SNP variation among 75 *Salmonella* genomes from multiple sources. Wild bird isolates from this study (green triangles) cluster within the genomic space predominantly occupied by poultry-associated strains. PC1 and PC2 account for 17.5% and 13% of the total genetic variance, respectively.

Consistent with the PCA results, SNP-based phylogenetic analysis showed that the isolates obtained in this study clustered predominantly with strains of poultry origin, including isolates derived from raw feed materials, transport materials, overshoes, and poultry farm environments (Fig. 3). Pairwise SNP distance analysis revealed low to moderate genomic divergence across sources (Fig. 4), with mean distances generally ≤10 SNPs among poultry-associated isolates and those from wild birds and humans. Notably, isolates from wild birds were closely related to broiler and human isolates (on average, ∼9.6 SNPs), indicating integration into a shared transmission network rather than forming a distinct lineage. Similarly, heavy breeder isolates displayed consistently low genetic distances (∼4.8 SNPs) across multiple sources, suggesting a central role in the dissemination of this lineage. In contrast, pig-derived isolates were markedly more divergent (24–33.6 SNPs), supporting the presence of a partially independent transmission cycle. The complete list of genomes included in the comparative phylogenetic analysis is provided in Supplementary Table S3.

**Figure 3.**
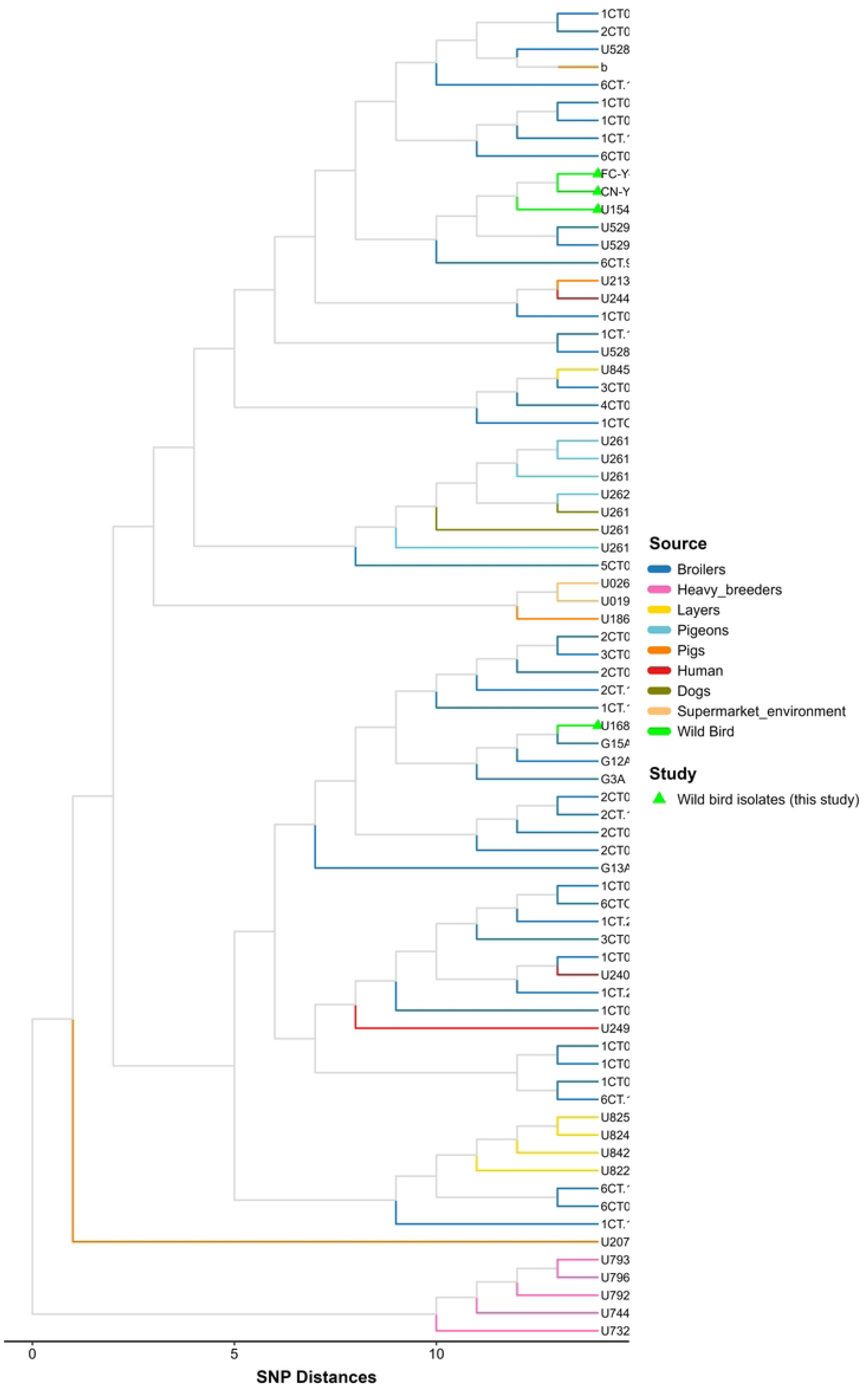
SNP-based phylogeny of *Salmonella* Infantis isolates from this study and diverse sources. Colored lines indicate an isolation source consistent with the PCA clustering.

**Figure 4.**
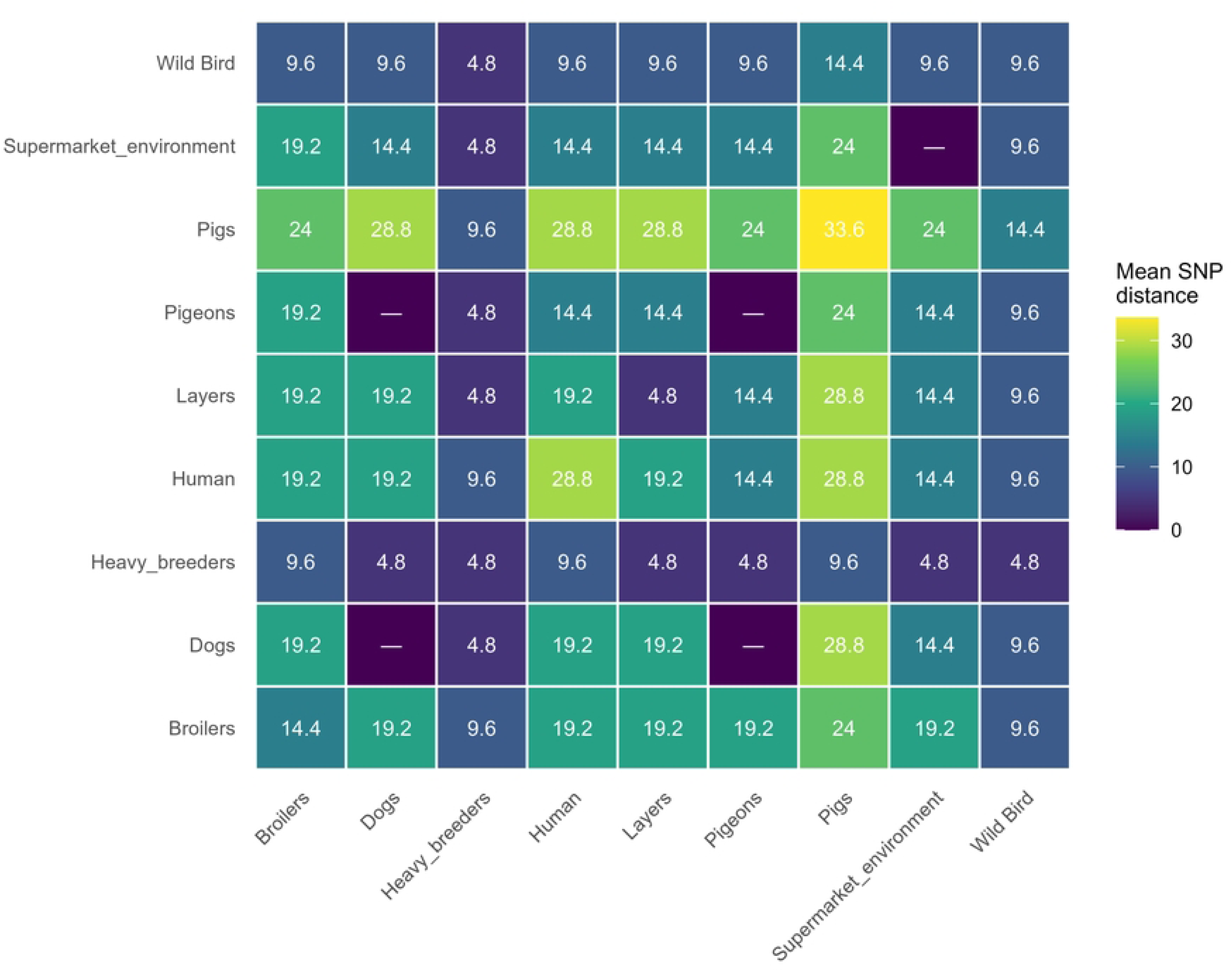
Mean pairwise Single Nucleotide Polymorphism (SNP) distance heatmap revealing genetic connectivity of *Salmonella* Infantis across human, animal, and environmental sources. **Notes**: Cells represent average SNP differences between sources. Lower values indicate higher genetic similarity. Most sources show low to moderate distances (≤10 SNPs), whereas pig-derived isolates are more divergent (24–33.6 SNPs)

## Discussion

The present study detected *Salmonella* in (3.73%) of fecal samples collected from two wild aquatic bird species, the Andean Coot (*F. ardesiaca*) and the Neotropical Cormorant (*P. brasilianus*) species of wild aquatic birds. Despite the low prevalence observed in this study, *Salmonella* has been previously reported in wild avian species, including *Anas platyrhynchos* [45], *Phalacrocorax carbo hanedae* [46], and other waterbirds [47,48]. These findings suggest that wild birds may become exposed to *Salmonella* in aquatic environments, and could contribute to its environmental dissemination across interconnected and agricultural ecosystems [49–52].

In wild birds, *Salmonella* infections are characterized by intermittent shedding, low host specificity, and a broad host range [9,53]. Salmonellosis caused by *Salmonella enterica* subsp. enterica serovar Infantis has been associated with sporadic mortality events, particularly among juvenile birds in breeding colonies or shared feeding areas where interspecies contact facilitates transmission [7,54]. These ecological characteristics may help explain the low prevalence observed in this study, as the sampled birds were clinically healthy adults inhabiting relatively extensive natural habitats where population density and interspecies contact are limited.

*Salmonella* Infantis was the predominant serotype identified in this study. Although this serovar is considered to have lower pathogenic potential in humans compared with some other *Salmonella* serovars, possibly due to reduced expression of genes associated with *Salmonella* pathogenicity island 1 (SPI-1) [55,56], it has become one of the most prevalent serovars reported in food-producing animals worldwide [57].

Multiple studies have linked *S.* Infantis with poultry production systems, including feed sources, broiler and layer farms, chicken carcasses, and on-farm environments such as manure, litter, and contact surfaces [58]. Additionally, recent studies have reported frequent detection of *Salmonella* in aquatic environments, with *S*. Newport, *S*. Infantis, and *S*. Typhimurium among the most prevalent serovars (Chen et al. 2024). These findings highlight the complex ecological interactions connecting agricultural systems, environmental reservoirs, and wildlife.

Our results demonstrated that *S.* Infantis isolates exhibited concordant phenotypic and genotypic antimicrobial resistance. The analysis revealed the presence of a pESI-like plasmid (PNUSAS047891) carrying multiple antimicrobial resistance genes (ARGs). Similar resistance patterns have been documented in aquatic and migratory wild birds [7,45,59]. These findings are particularly concerning; the broad application of these antibiotics in both human and veterinary medicine exacerbates the risk of disseminating clinically relevant resistance determinants [60–62].

In developing countries, the widespread availability of antimicrobials such as sulfonamides and trimethoprim has been associated with the dissemination of resistance genes, including *sul1* and *dfrA14* [63]. Similarly, β-lactams carrying with *blaCTX-M-65* and tetracyclines harboring *tet*(*A*) are commonly used in veterinary practices, often as growth promoters or prophylactic agents [61,64]. These practices contribute to the selection and maintenance of resistant bacterial populations, which can subsequently spread into natural ecosystems through the discharge of animal excreta and wastewater, particularly in areas where aquatic birds coexist with poultry and agricultural activities [65,66].

Genomic characterization of the isolates confirms that antimicrobial resistance in the *S*. Infantis isolates was associated with the presence of the pESI-like plasmid PNUSAS047891. This megaplasmid has been recognized as a key factor underlying the global success of *S*. Infantis because it carries a complex cluster of genes related to antimicrobial resistance, virulence, and tolerance to heavy metals and disinfectants [56]. Although the specific plasmid multilocus sequence type (pMLST) could not be determined due to locus identity limitations, its classification within incompatibility group IncI1 is consistent with the architecture of pESI-like plasmids commonly reported in intensive poultry production systems [67].

Principal Component Analysis (PCA) and SNP-based phylogenetic analyses demonstrate that the isolates cluster predominated with poultry-associated strains, reinforcing a poultry-linked origin. Specifically, the high genomic concordance between these isolates and those recovered from feed, transport matrices, and farm environments suggests the active dissemination of antimicrobial-resistant (AMR) bacteria from intensive production systems into the surrounding biosphere. These distribution patterns carry significant ecological implications, indicating potential spillover into wild bird populations and the contamination of shared habitats.

Notably, all Salmonella-positive samples were determined at Yahuarcocha Lake, which exhibits the highest phase of eutrophication and anthropogenic influence among the studied lakes. Previous studies have documented nutrient enrichment and water pollution associated with surrounding urban development, tourism, and agricultural activities [29,68]. Similar patterns for aquatic birds have been documented, suggesting that environmental degradation promotes pathogen circulation at wildlife-human interfaces [69]. In this context, local conditions likely enhance interspecific interactions among wildlife, domestic animals, and human-associated microbial sources, thereby increasing the risk of environmental exposure to enteric pathogens.

## Conclusion

The detection of Salmonella in wild aquatic birds highlights the potential role of wildlife as environmental sentinels of pathogen circulation and antimicrobial resistance in natural ecosystems. Genomic analyses revealed that the *Salmonella* Infantis isolates identified in this study are closely related to poultry-associated strains carrying pESI-like plasmids encoding multidrug resistance. These findings provide evidence of a link between antimicrobial-resistant bacteria from agricultural systems into surrounding environments, where wild birds may be exposed through contaminated water or food resources. Although direct transmission pathways remain difficult to determine, our results emphasize the importance of integrated environmental surveillance within a One Health framework.

## Data Availability Statement

All scripts and associated data are available at: https://github.com/Nivia-L/Salmonella_wild_birds.git. Genomic data will be deposited in a public repository upon acceptance.

## Acknowledgements

The authors thank Agusto Luzuriaga-Neira for the review and comments. The authors also thank to “DIRECCIÓN GENERAL DE INVESTIGATION (DGI)” for funding through Grant N°42. Whole-genome sequencing was supported by the U.S. Food and Drug Administration (FDA) of the U.S. Department of Health and Human Services (HHS) as part of a financial assistance award (5U19FD007122) with 100% funded by FDA/HHS. The contents are those of the authors and do not necessarily represent the official views of, nor an endorsement, by FDA/HHS or the U.S. government. The authors thank the UNIETAR-FMVZ staff for their assistance during the field laboratory, and the students Karla Mena and Leonardo Cedeño for their fieldwork assistance.

## Authorship contribution

N.R: conceptualization, data curation, formal analysis, investigation, methodology, writing-original draft; C.V: conceptualization, investigation, methodology, supervision, validation, writing-review & editing; J.M.: conceptualization, data curation, formal analysis, software, validation, writing - review & editing; M.L.I.: conceptualization, investigation, validation writing – review & editing; B.D.S.: conceptualization, investigation, validation writing – review & editing; D.A.: conceptualization, data curation, formal analysis, visualization, Writing – original draft; N.L.: conceptualization, data curation, formal analysis, funding acquisition, investigation, project administration, supervision, writing – review & editing **Authorship interest conflicts**: The authors declare no competing interests. Research

